# Superficial bound of the depth limit of 2-photon imaging in mouse brain

**DOI:** 10.1101/618454

**Authors:** Kevin Takasaki, Reza Abbasi-Asl, Jack Waters

## Abstract

2-photon fluorescence microscopy has been used extensively to probe the structure and functions of cells in living biological tissue. 2-photon excitation generates fluorescence from the focal plane, but also from outside the focal plane, with out-of-focus fluorescence increasing as the focus is pushed deeper into tissue. It has been suggested that the 2-photon depth limit, beyond which results become inaccurate, is where in- and out-of-focus fluorescence are equal. We found the depth limit of 2-photon excitation in mice with GCaMP6 indicator expression in all layers of visual cortex, by comparing near-simultaneous 2- and 3-photon excitation. 2-photon results were accurate only superficial to 450 μm, matching the depth at which in-and out-of-focus fluorescence were equal. The expected depth limit is deeper in tissue with fewer fluorophores outside the plane of interest. Our results, from tissue with a largely homogenous distribution of fluorophores, establish a superficial bound on the 2-photon depth limit in the mouse visual cortex.

## INTRODUCTION

2-photon excitation permits fluorescence imaging with cellular and subcellular resolution hundreds of micrometers into biological tissue. Generally, the maximal imaging depth (depth limit) of 2-photon excitation is determined by fluorescence from outside the focal plane. As the focal plane is pushed deeper into scattering tissue, illumination intensity at the tissue surface must be increased to maintain intensity in the focal plane, resulting in an increase in out-of-focus fluorescence with increasing depth (Ying *et al.*, 1999; Theer *et al.*, 2003). In a seminal study, Theer and Denk (2006) explored 2-photon excitation analytically and defined the fundamental imaging depth limit by calculating the depth at which the detected fluorescence generated by ballistic and scattered excitation light outside the focal plane equals that from fluorophores excited in the ballistic focus. The ratio of in- and out-of-focus fluorescence is a complex function of numerous factors, including numerical aperture, laser pulse duration, scattering anisotropy, and fluorophore distribution, but the calculations of Theer and Denk (2006) suggest that the depth limit is at ~3 scattering length constants under typical imaging conditions. In cortical grey matter, 3 scattering length constants corresponds to ~600 μm below the tissue surface.

3-photon excitation permits deeper imaging than 2-photon excitation, in part because 3-photon excitation generates fluorescence almost exclusively from the focal plane (Horton *et al.*, 2013; Kobat *et al.*, 2009; Kobat *et al.*, 2011; Ouzounov *et al.*, 2017; Yildirim *et al.*, 2019). In the absence of out-of-focus fluorescence one expects the functional properties of neurons measured with 2- and 3-photon excitation to be identical, but this expectation has not been tested directly and the impact of out-of-focus fluorescence has not been measured. 3-photon excitation offers the opportunity to estimate in- and out-of-focus fluorescence and thereby test the predictions of earlier analyses. We implemented near-simultaneous 2- and 3-photon excitation to compare results 200 to 650 μm below the surface of the brain in transgenic mice with dense GCaMP6 expression throughout neocortex. Our results show that 2- and 3-photon excitation produce equivalent results in superficial layers but not in deep in cortex, and indicate that the depth limit of 2-photon excitation corresponds to the plane where in- and out-of-focus fluorescence are equal, consistent with Theer and Denk (2006).

## RESULTS

As an illumination source for 3-photon excitation, we used a 40 Watt Coherent Monaco laser source and Opera-F optical parametric amplifier, providing 2 μJ, 50 fs pulses at 1 MHz. We configured a MIMMS 2-photon microscope for 3-photon excitation, exchanging the scan and tube lens to increase transmission through the microscope at 1300 nm and added a compressor to compensate for pulse dispersion between the laser source and sample. Through a cranial window over visual cortex, we were routinely able to image neurons >1 mm below the pial surface of cortex in GCaMP6 mice (supplementary figure 1). Fluorescence intensity followed a cubic relationship with illumination intensity, consistent with fluorescence being driven by the absorption of 3 photons.

In mice expressing GCaMP broadly in cortical pyramidal neurons, loss of contrast was noticeable in 2-photon images from hundreds of micrometers below the brain surface (figure 1A) where contrast was preserved by 3-photon excitation (figure 1B). To compare 2- and 3-photon excitation more directly, we implemented near-simultaneous 2- and 3-photon excitation. We used 2 laser sources, combining the beams immediately before the scanning galvanometers (figure 2A). With a fast Pockels cell on each laser line acting as a shutter, we alternated 2- and 3-photon excitation, line-by-line (figure 2B). The line duration was 0.5 ms, resulting in 0.5 ms separation of 2- and 3-photon images.

**Figure 1.**
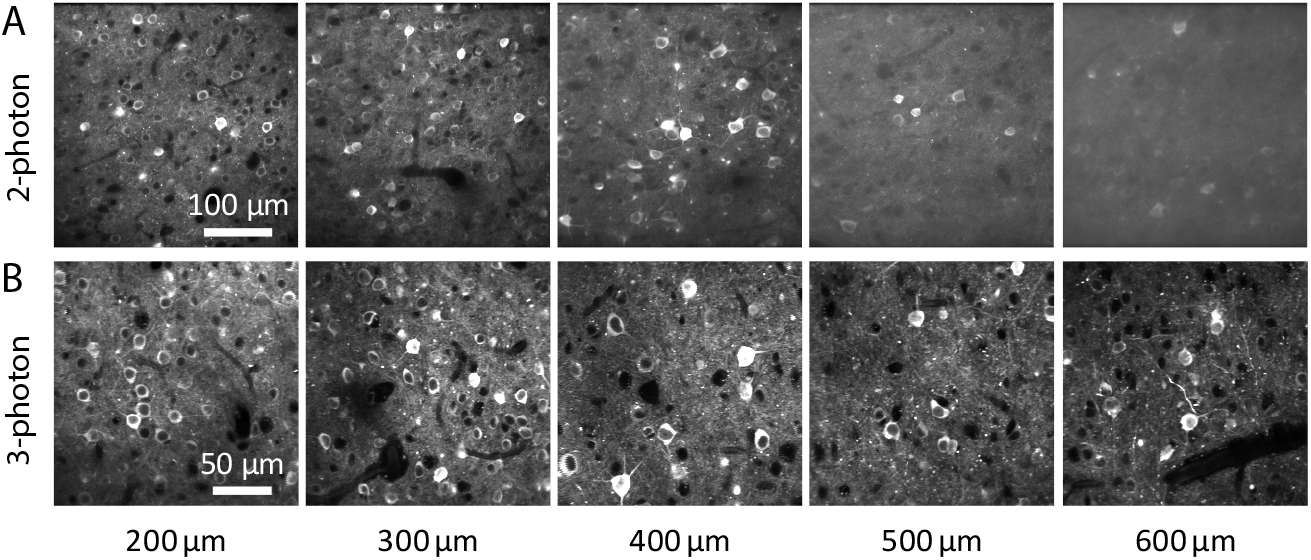
Contrast declines with depth with 2-photon excitation. (A) Images acquired using 2-photon excitation, focused 200-600 μm below the pial surface of visual cortex. Emx1-IRES-Cre;CaMK2a-tTA;Ai94 mouse (B) Images acquired from the same mouse using 3-photon excitation. 2- and 3-photon images are from different fields of view.

**Figure 2.**
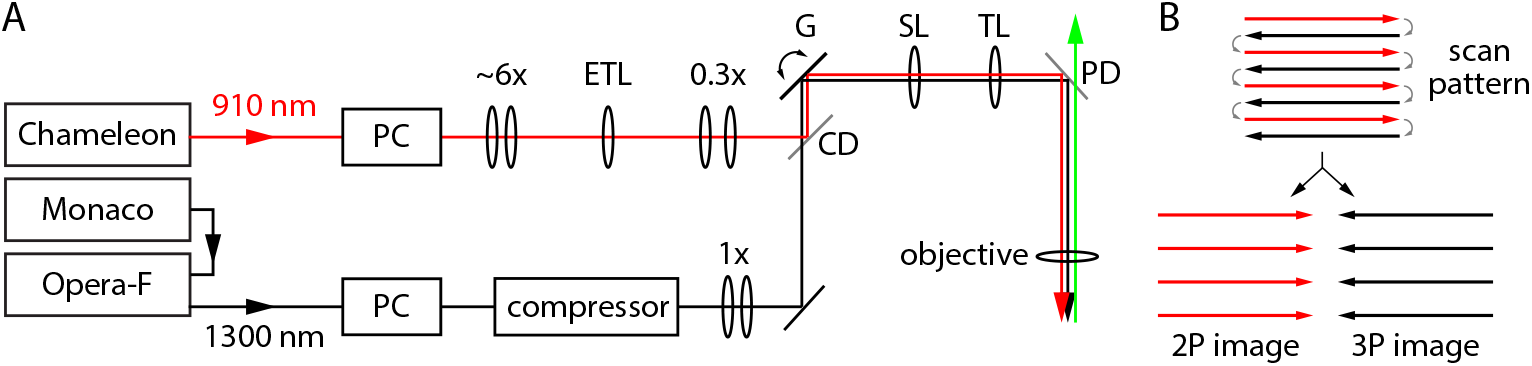
Implementation of near-simultaneous 2- and 3-photon excitation. (A) Schematic of the optical layout for near-simultaneous 2- and 3-photon excitation. 1300 nm beam (black) passed a Pockels cell [PC], prism compressor, a collimating telescope, combining dichroic mirror [CD], x-y galvanometer pair [G], scan lens [SL], tube lens [TL], FF735-DI02 primary dichroic mirror [PD] and objective lens. 910 nm beam (red) passed a Pockels cell (PC), beam expansion to ~1 cm diameter, electrically-tunable lens [ETL], 0.3x beam expansion before being reflected by the combining dichroic mirror onto the galvanometer pair. (B) Scanning strategy for near-simultaneous 2P- and 3-photon excitation. Red: 920 nm excitation, no 1300 nm excitation. Black: no 920 nm excitation, 1300 nm excitation. Grey: both lasers blocked. Lines were sorted into 2- and 3-photon images.

In superficial cortex, 2- and 3-photon results were similar. The same neurons were visible in near-simultaneous 2- and 3-photon images and changes in fluorescence were coincident in 2- and 3-photon image pairs (supplementary movie 1); the results of motion correction and segmentation on 2- and 3- photon movies were similar (standard deviations of motion correction distributions <2 μm at <350 μm, figure 3C); there were 50-90 neurons identified in each image (figure 3D); >80% of neurons in 3-photon images matched a neuron in the corresponding 2-photon image (figure 3E); and traces extracted from matching neurons in 2- and 3-photon movies were strongly correlated, with Pearson correlation coefficients of ~0.8-0.9 (figure 3F), consistent with a previous study (Wang *et al.*, 2017).

**Figure 3.**
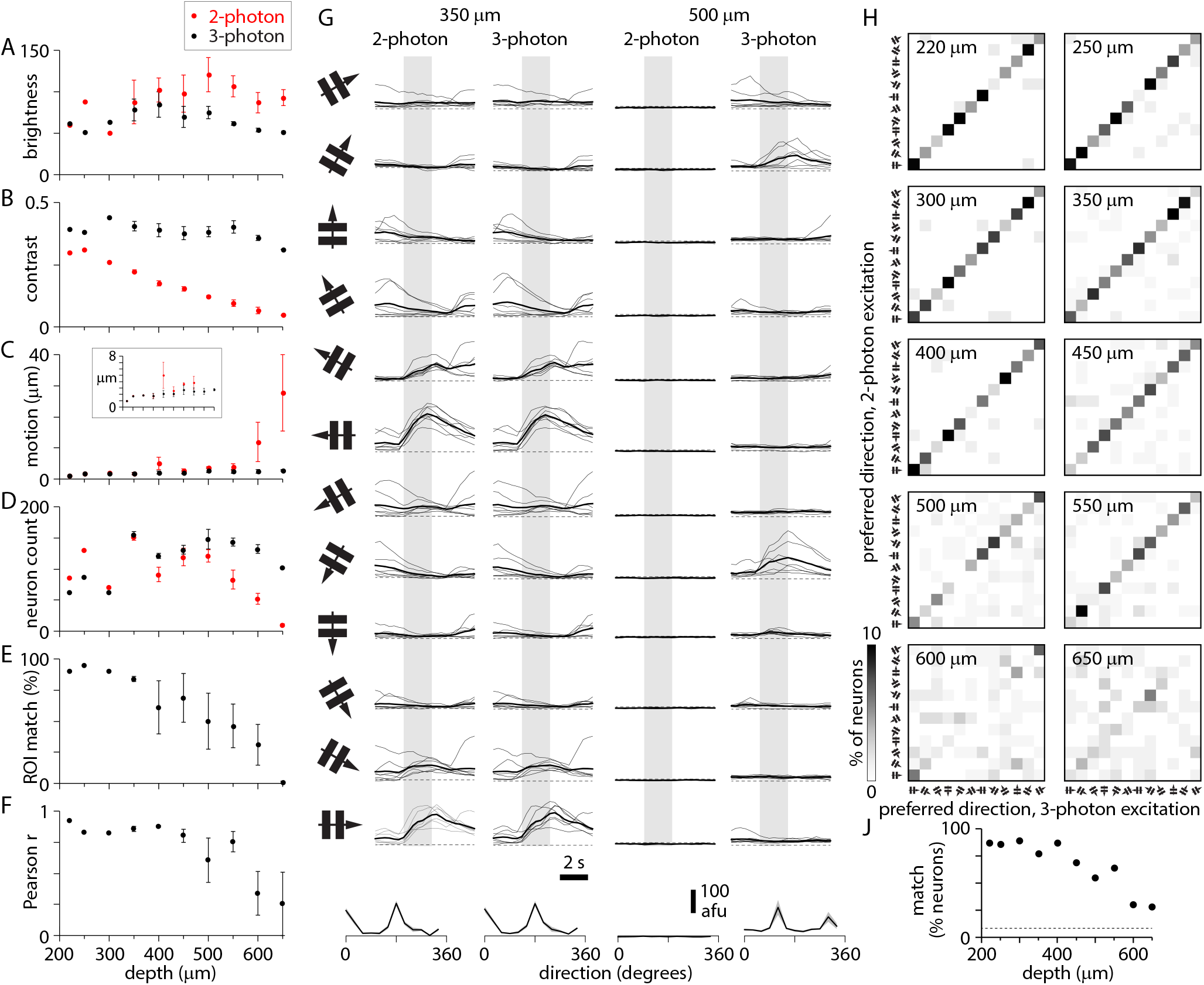
Changes in 2-photon image quality and apparent ΔF with depth. (A-D) Plots of image brightness (A), contrast (B), corrected motion (C) and ROI count(D) for 2-photon (red) and 3-photon excitation (black), plot as a function of depth below the pial surface of cortex. Mean ± SEM of 3 experiments from 2 Slc17a7-Cre;Ai162 mice. (E) ROI match (percentage of 3-photon ROIs also segmented from 2-photon images) as a function of depth. (F) Pearson correlation coefficient between 2- and 3-photon fluorescence traces, plot as a function of depth. (G) 2- and 3-photon changes in fluorescence to grating stimuli for two neurons, 350 and 500 μm below the pia. Each panel shows change in fluorescence (in arbitrary fluorescence units) through time during presentation of the drifting grating (icon to left indicates orientation and direction) for 2 seconds (grey bar). 8 individual traces and the mean (thick line) per direction. Dashed line indicates zero fluorescence. Below: resulting direction tuning curve. (H) Plots comparing preferred direction of neurons measured with 2-photon (y axis) and 3-photon (x axis) excitation, for each depth. Colors indicate percentages of the total number of neurons at each depth (zero is white, 10% is black, see color bar). Directions progress at 30 degree intervals from the low left corner of each plot (icons). (J) Percentage of neurons with matching direction preferences measured with 2- and 3-photon excitation, from 200 to 650 μm. Dashed line: 8.3%.

The similarity of 2- and 3-photon results declined with depth. In 3-photon images, image contrast, motion correction, and number of neurons changed little with depth. In 2-photon images, contrast declined incrementally with depth, to near zero at 650 μm (figure 3B). Lateral motion correction from 2-photon movies increased with depth: the standard deviation of motion correction was <3 μm at <400 μm; at 650 μm, the standard deviation of lateral motion correction was <3 μm for 3-photon excitation and ~25 μm for 2-photon excitation (figure 3C, supplementary figure 1). The segmentation routine identified few neurons in deep locations (figure 3D), and the overlap between matching neurons in 2- and 3-photon images and the correlation coefficient between the resulting traces both declined at >350-400 μm (figure 3E and F).

To determine how the decline in image quality with depth affects the functional properties of cortical neurons measured with 2-photon excitation, we examined the apparent responses of cortical neurons to visual stimuli. We presented sinusoidal gratings drifting in 12 directions, and calculated the direction preference of each neuron from extracted fluorescence traces, comparing results from 2- and 3-photon excitation. For superficial neurons, visually-evoked changes in 2- and 3-photon fluorescence were almost identical, trial-by-trial (figure 3G) and the resulting preferred direction of each neuron was closely matched (figure 3H), with 83% (305 of 368) of neurons ≤350 μm from the brain surface exhibiting identical preferred directions with 2- and 3-photon excitation. Visually-evoked changes in 2-photon fluorescence were suppressed in deeper neurons (figure 3G, supplementary figure 3) and the percentage of neurons with matching 2- and 3-photon direction preference declined (figure 3H & J, supplementary figure 4). At 600 μm, the number of neurons with matching preference was above chance (1/12 = 8.3%), but <<50 %.

2-and 3-photon excitation produce equivalent results from superficial depths, but the results become less similar >400 μm below the brain surface. Increasing out-of-focus fluorescence and the resulting decline in image contrast are the likely cause. From the ratio of contrast in 2- and 3-photon images, we estimated the percentage of fluorescence that originated from the focal plane during 2-photon excitation. As expected, the percentage of 2-photon fluorescence originating from the focal plane decreased with increasing depth (figure 4A, supplementary figure 6). In- and out-of-focus fluorescence were equal at ~400-450 μm, the depth beyond which the results of 2-photon excitation are inaccurate. Hence our results support the depth limit corresponding to the depth at which in- and out-of-focus fluorescence are equal.

**Figure 4.**
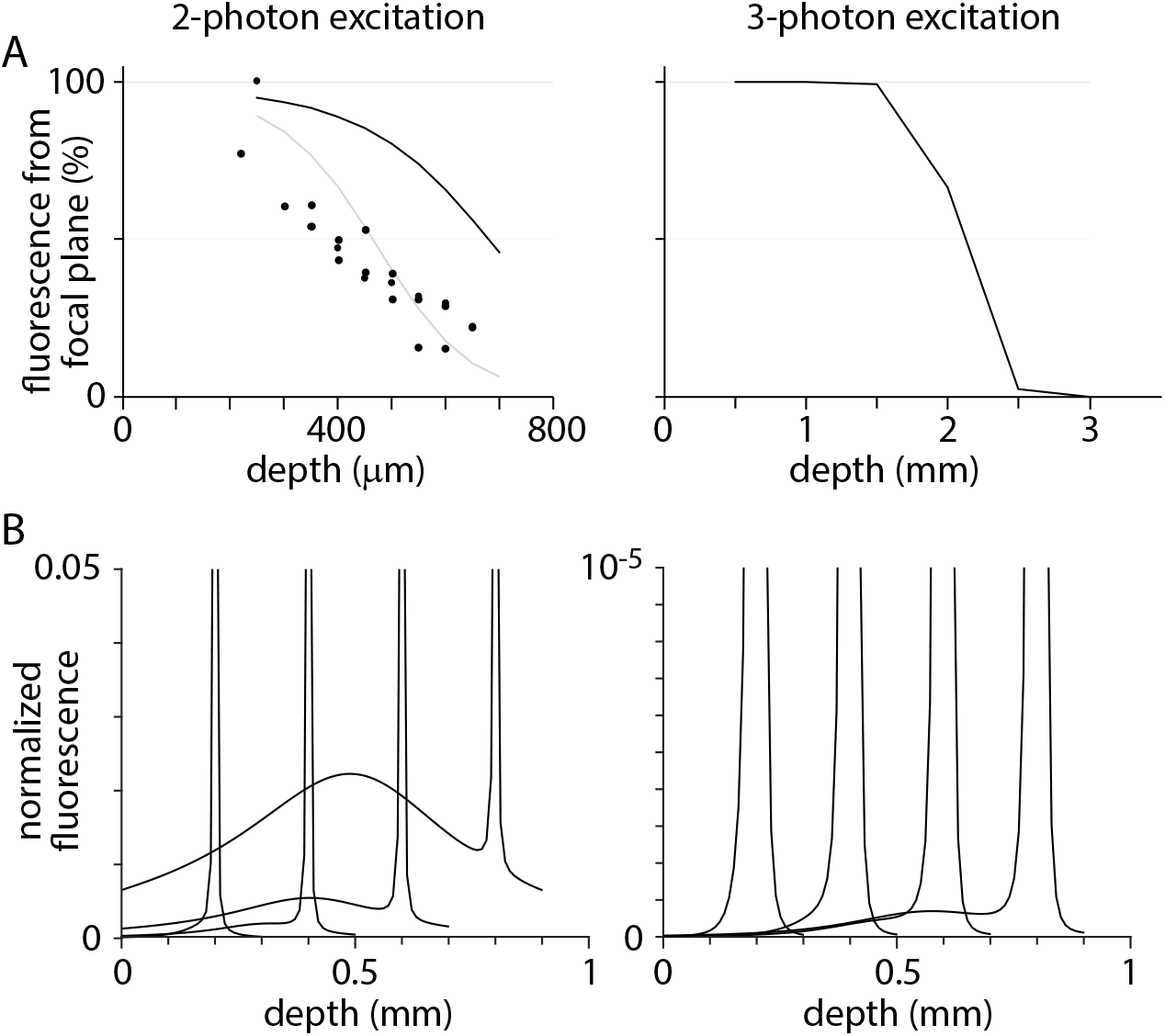
In- and out-of-focus fluorescence. (A) Percentage of total fluorescence that originates from the focal plane, plot as a function of depth of the focal plane below the brain surface. Each point represents a single measurement (from a movie at one depth in one mouse). Lines are calculated from equation 1 with scattering length constants of 200 μm (black) and 150 μm (grey). (B) Plots showing the depth from which fluorescence originates with the focal plane at 200, 400, 600 and 800 μm below the brain surface. Fluorescence was calculated with equation 1 and normalized to that in the focal plane. Note the difference in scale for 2- and 3-photon excitation. Breakdown of fluorescence sources in supplementary figure 7.

We compared our measurements of in- and out-of-focus fluorescence with predictions from theoretical modeling of focused light propagation in scattering tissue (Ying, et al. 1999; Theer & Denk, 2006;). According to this model, 50% in-focus fluorescence occurs ~3 scattering lengths below the brain surface, ~450 μm for a scattering length of 150 μm and 600-700 μm for a scattering length of 200 μm (figure 4A, black and grey lines, respectively). 3-photon excitation is almost free of out-of-focus fluorescence at these depths.

2-photon excitation will support imaging >450 μm below the brain surface if there are few fluorophore molecules outside the focal plane. Unfortunately, out-of-focus fluorescence arises from fluorophores throughout the tissue above and, to a lesser extent, below the focal plane (figure 4B; Theer & Denk, 2006). 3-photon excitation generates little out-of-focus fluorescence, but again most arises from locations immediately superficial to the focal plane. Hence a substantial reduction in out-of-focus fluorescence, and increase in depth limit, would likely occur only in tissues with few fluorophore molecules throughout the entire depth of tissue above and below the focal plane.

## DISCUSSION

We compared the results of 2- and 3-photon excitation of GCaMP6s in excitatory neurons in mouse visual cortex. Results from superficial cortex were similar, an expected result that confirms that the cellular signals reported by GCaMP6s are independent of the mechanism of excitation, and that 3P imaging has not been compromised by saturation or phototoxic effects (Yildirim, et al. 2019). With increasing depth from ~250-650 μm, 2-photon image contrast declined and 3-photon image contrast was preserved. Many measures (estimated motion, number of neurons segmented, matching of segmented neurons, correlation traces, similarity of fluorescence changes, similarity of preferred direction) were robust to changes in 2-photon image contrast to ~400 μm, but deteriorated between 400 and 550 μm on average, some abruptly, compromising measurement of fluorescence changes and direction tuning.

In our experiments, we used a mouse line with GCaMP6s expression in excitatory neurons through all layers of cortex. From the perspective of out-of-focus fluorescence, we expect these mice to be a worst-case scenario for 2-photon excitation. In these mice, our results place the depth limit at ~450 μm below the brain surface, shallower than the depth predicted Theer and Denk (2006) and by our calculations. We expect aberrations to reduce the depth at which in- and out-of-focus fluorescence are equal. Aberrations are present in any imaging system, but not included in our calculations or those of Theer & Denk (2006). Slight compression of cortex is common in cranial window preparations (e.g. de Vries *et al.*, 2018) and might further reduce the depth limit by reducing the scattering length of cortical tissue. Hence one expects the depth limit of 2-photon excitation to be shallower than suggested by calculations. Our measurements indicate that the depth limit can be as shallow as 2-3 scattering lengths or ~450 μm.

Our results drive two predictions that we have not tested directly. Firstly, we expect that 2-photon excitation will be adequate for characterization of functional properties such as direction tuning in neurons ≤450 μm from the brain surface in nearly all GCaMP6s mouse lines. Secondly, we expect 2- and 3-photon results to be comparable at >450 μm in many preparations. We observed substantial mouse-to-mouse variability at 500-650 μm, suggesting that 2-photon excitation might be a viable tool to >450 μm in a small subset of our mice. In other mouse lines and tissues, 2-photon excitation at >450 μm will provide more accurate functional measurements in preparations with less out-of-focus fluorescence, including tissues with sparser expression of GCaMP6s and tissues labeled with indicators with low resting fluorescence, such as jGCaMP7c (Dana *et al.*, 2018). In such tissues, there are several factors that might limit 2-photon excitation. Out-of-focus fluorescence, though reduced, will still occur and may equal in-focus fluorescence at a location deeper than 450 μm. Aberrations might prove limiting, enabling adaptive optics to extend the depth limit (Ji *et al.*, 2010; Ji *et al.*, 2012). A third possibility is maintenance of image quality to a depth at which the thermal limit of brain tissue is met (Podgorski & Ranganathan, 2016).

In summary, we have established that 2- and 3-photon excitation are equivalent ≤~450 μm below the brain surface in mice with GCaMP6s expression throughout cortical layers. Tentatively, we suggest the depth limit of 2-photon excitation is 450 μm or deeper in nearly all mouse lines, since few if any mice express a higher proportion of fluorophore molecules outside the focal plane than mice with expression throughout the cortical layers. In tissues with and tissues without extensive fluorophore expression outside the focal plane, 3-photon excitation enables measurement of cellular activity beyond the depth limit of 2-photon excitation.

## METHODS

### Basic 3-photon microscope

Our 3-photon microscope was built around a Coherent Monaco / Opera-F laser source (≤2 nJ, 50 fs pulses at 1 MHz; Coherent Inc., Santa Clara) and a modified MIMMS microscope manufactured by Sutter Instrument (Novato, CA). We replaced the scan and tube lenses (respectively, Thorlabs SL50-3P and a Plössl pair of achromatic doublets, Thorlabs AC254-400-C) to improve transmission at 1300 nm. The primary dichroic mirror was FF735-DI02 (Semrock, Rochester NY). We used an Olympus 25x/1.05 objective (75% transmission at 1300 nm) or Nikon 16×/0.8 objective (50% transmission at 1300 nm) and image acquisition was controlled by ScanImage (Vidrio Technologies LLC) with acquisition gating for low rep rate lasers.

We estimated group delay dispersion (GDD) through the microscope at ~15,000 fs^2^, approximately half of which was attributable to the Pockels cell (360-40-03-LTA, Conoptics Inc, Danbury CT). To compensate, we built a 4-pass pulse compressor using a single SF-11 glass prism (Thorlabs PS-853) and a two hollow roof prism mirrors (Thorlabs HRS1015-P01 and HR1015-P01). Compression was tuned by maximizing brightness with a fluorescein sample.

400-500 mW of 1300 nm illumination was available after the objective, corresponding to transmission from laser source to sample of ~20%. The maximum field of view of 3-photon excitation was 360 x 360 μm. Images were acquired with dual linear galvanometers at a frame rate of ~8 Hz.

### Illumination intensity

Photodamage is often a concern in light microscopy. Photodamage can result from linear processes, principally heating (resulting from the absorption of infrared light by water in brain tissue) and from non-linear processes. Non-linear processes are of particular concern with high-energy pulsed sources such as those used for 2- and 3-photon fluorescence microscopy. Heating-related photodamage often occurs with >250 mW of prolonged illumination at 800-1040 nm (Podgorski & Ranganathan, 2016) and the molar extinction coefficient of water at 1300 nm is ~2x that at 900 nm (Curcio & Petty, 1951; Hale & Querry, 1973; Bertie & Lan, 1996), suggesting that heating-related tissue damage may occur at >~100-150 mW of prolonged illumination at 1300 nm. To avoid damage, we used illumination intensities <100 mW. Typically, we could image through the depth of neocortex using <30 mW illumination while maintaining a signal-to-noise ratio comparable to typical 2-photon experiments. We rarely observed signs of photodamage, even in mice subjected to 2 hours of continuous 3-photon imaging per day for 5 days.

### Near-simultaneous 2- and 3-photon excitation

For 2-photon excitation, we used a Coherent Chameleon Ultra II laser source at 920 nm. For near-simultaneous 2- and 3-photon excitation, we used a Nikon 16×/0.8 objective (50% transmission at 1300 nm). Time-averaged power available after the objective was 200-250 mW at 1300 nm. To match the focal planes of 2- and 3-photon excitation, to the 2-photon path we added an electrically-tunable lens (EL-10-30-TC, Optotune, Dietikon, Switzerland).

### Mice and surgeries

We used Cre-lox transgenic mice to drive GCaMP6s expression in excitatory neurons throughout cortical layers and areas, crossing Emx1-IRES-Cre *(B6.129S2-Emxltml(cre)Krj/J,* JAX stock number 005628; Gorski *et al.*, 2002) or Slc17a7-IRS2-Cre *(B6;129S-Slc17a7^tm11(cre)Hze^/J,* JAX stock number 023527; Harris *et al.*, 2014) and Ai162(TIT2L-GCaMP6s-ICL-tTA2 reporter mice (JAX stock number 031562, Daigle *et al.*, 2018).

A chronic cranial window was implanted over visual cortex as described previously (Goldey *et al.*, 2014; de Vries *et al.*, 2018). Briefly, under 0.5-2% isoflurane anesthesia, a head restraint bar was attached to the skull using C & B Metabond (Parkell) and a 5 mm diameter craniotomy was opened over the left visual cortex at coordinates 2.7 mm lateral, 1.3 mm anterior to lambda. A durotomy was performed and the craniotomy was sealed with a stack of three #1 coverslips, attached to each other using optical adhesive, and attached to the skull with Metabond.

### Visual stimuli

Visual stimuli were full-field sinusoidal gratings of 6 orientations, each drifting perpendicular to its orientation (12 directions), at spatial frequencies of 0.04 and 0.08 cycles per degree and a temporal frequency of 1 Hz. Each grating was presented 8 times in random order, each for 2 seconds with 1 second of grey screen between presentations. 0 degrees corresponds to a grating drifting horizontally in the nasal-to-temporal direction and 90 degrees to a downward-drifting grating. The visual stimulus display and its calibration were as described previously (de Vries *et al.*, 2018). Briefly, stimuli were displayed on an LCD monitor, 15 cm from the right eye, gamma-corrected and of mean luminance of 50 cd/m^2^. Spherical warping was employed to ensure the apparent size, speed, and spatial frequency were constant across the monitor.

### Image analysis

Image analysis was performed using custom routines in Python 3. For comparison of 2- and 3-photon excitation, images were first separated into 2- and 3-photon movies. Dark current, the mean of several images acquired with no laser illumination, was measured in each movie and subtracted. Image brightness (figure 4A) was measured in digitizer units. To avoid artifacts, each movie was normalized to the same mean brightness.

Image contrast (figure 4B) was expressed on a scale from 0 (no contrast) to 1. Contrast was calculated locally (in 22.5 x 22.5 pixel blocks) from the temporal mean projection of a movie, the final value being the mean of all the blocks. Contrast in each block was defined as 1 – minimum brightness / maximum brightness.

Each movie was motion-corrected and putative neuronal somata identified by segmentation. Soma and neuropil fluorescence traces were extracted and neuropil fluorescence was subtracted from the corresponding soma trace (r = 1). Motion correction, segmentation and trace extraction were performed using Suite2p (Pachitariu *et al.*, 2017) with default settings except for maxregshift which was set to 0.2 to permit ≤~70 μm motion correction in each transverse axis. Motion correction (figure 4C) was the mean of x- and y-corrections applied by Suite2p. Neuron count (figure 4D) was the number of putative somata returned by Suite2p, with manual editing to assist the sorting of somatic from non-somatic regions of interest. % match (figure 4E) was the percentage of putative neurons segmented in the 3- photon image that were also segmented in the corresponding 2-photon image, assessed manually by comparing images of segmented regions. Pearson correlation coefficient (figure 4F) was calculated from the neuropil-subtracted fluorescence traces using scipy.stats.pearsonr. To ensure that the correlation coefficient calculation was from matching regions of interest, traces were extracted from 2-photon and 3-photon movies using regions of interest segmented from 3-photon movies.

To compare 2- and 3-photon measurements of responses to drifting gratings, we used two measures: mean fluorescence response and preferred direction. Again, these measures were applied to traces extracted from 2- and 3-photon movies using regions of interest segmented from 3-photon movies. For each measure, we first calculated the mean response of each neuron (from 8 presentations). For the mean fluorescence response, we plot 2- vs 3-photon amplitudes of the mean change in fluorescence for each grating. Hence in the mean fluorescence response plots (figure 5A), each neuron is represented by 24 data points (12 directions x 2 temporal frequencies). The direction preference plots (figure 5B), report the grating direction that evoked the largest change in fluorescence and each neuron is therefore represented by a single data point.

### Modeling in- and out-of-focus fluorescence

To estimate the out-of-focus fluorescence generated by excitation light focusing through a homogeneous volume of fluorescent and scattering tissue, we modeled the intensity of ballistic and scattered light, *I_b_* (*z, ρ*) and *I_s_* (*z, ρ*) respectively, in a plane transverse to the optical axis defined by the polar radius, *ρ*, and depth *z* below the surface of the brain. We calculated the out-of-focus, 2-photon-excited fluorescence (F_oof_) numerically, following Theer & Denk (2006).

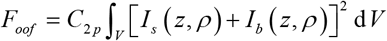

where, *V* is the out-of-focus illuminated volume of tissue, *C*_2*p*_ is a modality-specific scaling factor incorporating contributions from fluorophore concentration and excitation efficiency, and assumed to be constant over the volume.

We neglected possible depth dependence of fluorescence collection and detection, non-conservative attenuation due to bulk absorption of near-IR light, and the time dependence of excitation by ultrashort pulses that becomes a significant factor for pulse widths < ~50 fs (Theer & Denk, 2006; but see also Leray *et al.*, 2007).

Previous models (Xu & Webb, 1996; Theer & Denk, 2006) neglected the difference in distances traveled through tissue by on-axis and marginal rays. The difference in distance can be substantial for high-numerical aperture objectives, but of marginal importance when the focal plane is many multiples of the scattering length below the tissue surface. Here, we calculated fluorescence with the focal plane 1-4 scattering lengths below the tissue surface and therefore account for the dependence on propagation angle relative to the optical axis by incorporating a radially varying propagation distance,

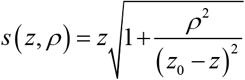

where *z*_0_ is the focal plane depth. This factor modifies the intensity profiles of *I_b_* and of *I_s_*.

*F_oof_* can be decomposed into individual contributions from ballistic, scattered, and cross-term interaction excitation, for 2-photon excitation:

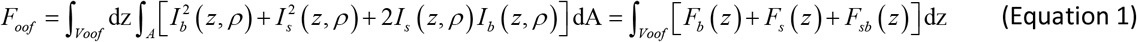

where, *V_oof_* is the out-of-focus volume denoting the range 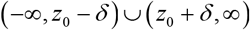 *δ* is the exclusion depth of in-focus light around *z*_0_.

In our calculations, *δ* was a fifth of the scattering length, or 40 μm, which we assume to be larger than the depth of focus and therefore underestimates the magnitude of the background; wavelength was 900 nm; numerical aperture 0.8; and anisotropy factor 0.9.

To calculate ballistic and scattered light intensities, we considered a Gaussian beam propagating from the surface (*z* = 0) of a scattering medium of scattering length 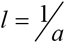 to a ballistic focus located at *z* = *z*_0_. We introduced a direction dependent propagation length 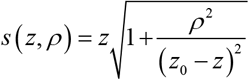, and calculate the ballistic intensity profile at depth *z* and radial distance *ω* according to

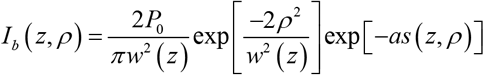

where, 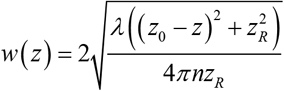 is the 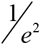 width 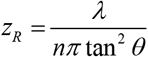 is the Rayleigh length determined by the NA-derived focusing half-angle.

As in Theer & Denk (2006), we calculated the intensity distribution of scattered light from a beam spread function derived for small-angle scattering (McLean, Freeman & Walker, 1998). We integrated over temporal and angular coordinates to obtain the normalized spatial distribution function

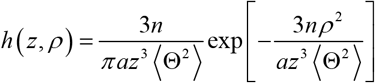

The spreading parameter 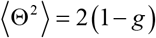 is derived from the anisotropy factor *g* and the function *h*(*z,p*) accounts for the diffusive spreading of scattered light with increasing depth from an initial on-axis ray, with total power increasing with depth according to 1 – exp [-*az*], modeling the transfer ofenergy from the ballistic to the scattered field.

Integrating over the initial surface distribution, the intensity distribution of scattered light at depth *z* is

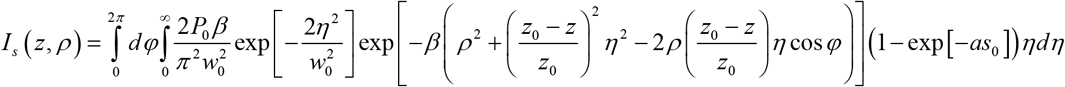

where 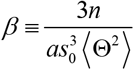.

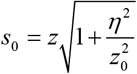 is the propagation distance from the surface

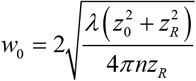 is the Gaussian beam width at the surface.

### Proportion of fluorescence originating from the focal plane

Calculation of the ratio of in- and out-of-focus fluorescence was based on image contrast. We subdivided the 256×512 pixel images of the motion-corrected, time-averaged 2-photon and 3-photon movies into 32×32 pixel subregions. Within each subregion, we determined the minimum pixel value and pixel value mean, 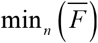 and 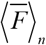 respectively, where 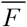 denotes the time-averaged fluorescence in each pixel with the minimum and mean functions over the 32×32 = 1024 pixels. For each subregion in each imaging modality (2P and 3P), we then calculated a contrast parameter, 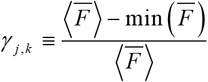, for the *j*-th subregion in the *k* = {2,3} (2P,3P) modality.

To calculate in- and out-of-focus fluorescence, we made three assumptions. Firstly, we assumed the time-averaged fluorescence in each pixel reflects the sum of the in-focus and out-of-focus fluorescence 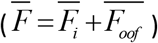. Secondly, we assumed 3-photon excitation generates no out-of-focus fluorescence so that 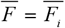 for 3-photon excitation. Thirdly, we assumed in-focus fluorescence is proportional to a modality-independent concentration factor, *C*, with a modality-dependent proportionality constant, so that *F_i_k__* = *ρ_k_C*.

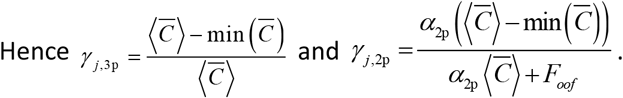

As a measure of the percentage of fluorescence that originates from the focal plane, we calculated an empirical contrast ratio (ECR): 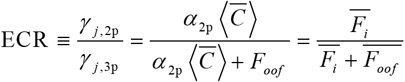

The ECR calculated in each subregion was averaged over the subregions to determine the time-averaged ECR for a given imaging depth.

We calculated the theoretical contrast ratio via a signal-to-background ratio calculation. We modeled the total in-focus fluorescence, *F_i_*, according to 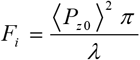 where *P*_*z*0_ is the total, scattering attenuated, ballistic power estimated at the focal plane according to 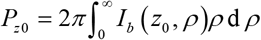. The signal-to-background ratio was defined as the ratio of total in-focus to out-of-focus fluorescence, given by 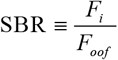 which ranges from 0 at very large depths to ∞ in the background-free case. We defined the contrast ratio, 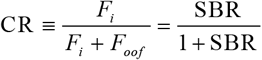 to range from 0 to 1.

## ACKNOWLEDGEMENTS

We thank the Allen Institute founder, Paul G. Allen, for his vision, encouragement and support. We thank members of the Research Engineering and Neural Coding teams for helpful discussions and Ariel Leon for hardware support.

---

**Supplementary movie 1. Examples of simultaneous 2- and 3-photon image pairs at different depths**. Examples of matched 2- and 3-photon movies 250, 450 and 650 um below the pia. 2- and 3-photon movie pairs were acquired pseudo-simultaneously. Each movie was acquired at a different illumination intensity and each was scaled differently for display purposes. Slc17a7-Cre;Ai162 mouse.

**Supplementary figure 1.**
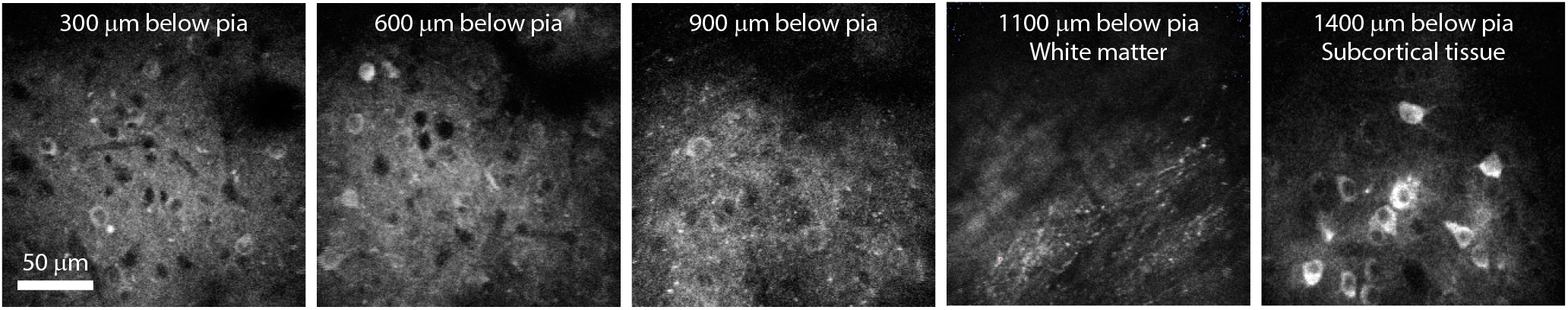
Deep imaging of GCaMP6s fluorescence using 3-photon excitation. Example 3-photon images from 300, 600, 900, 1100 and 1400 μm below the pial surface of visual cortex. Emx1-IRES-Cre;CaMK2a-tTA;Ai94 mouse.

**Supplementary figure 2.**
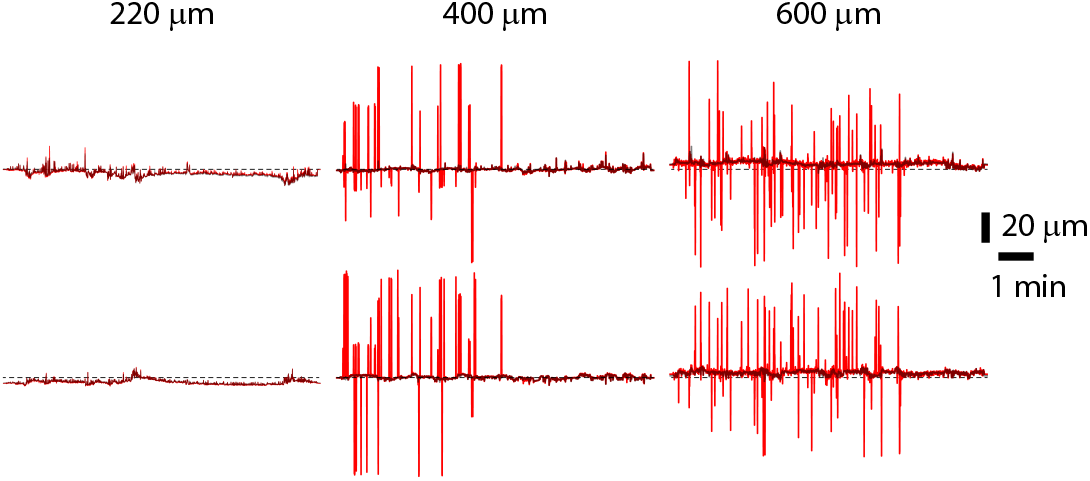
Examples of motion. An example of motion estimates (in μm) from the motion correction routine, for three depths. Two plots per depth for translations in the two transverse dimensions, relative to the optical axis. Red, 2-photon estimated motion; grey, 3-photon estimated motion.

**Supplementary figure 3.**
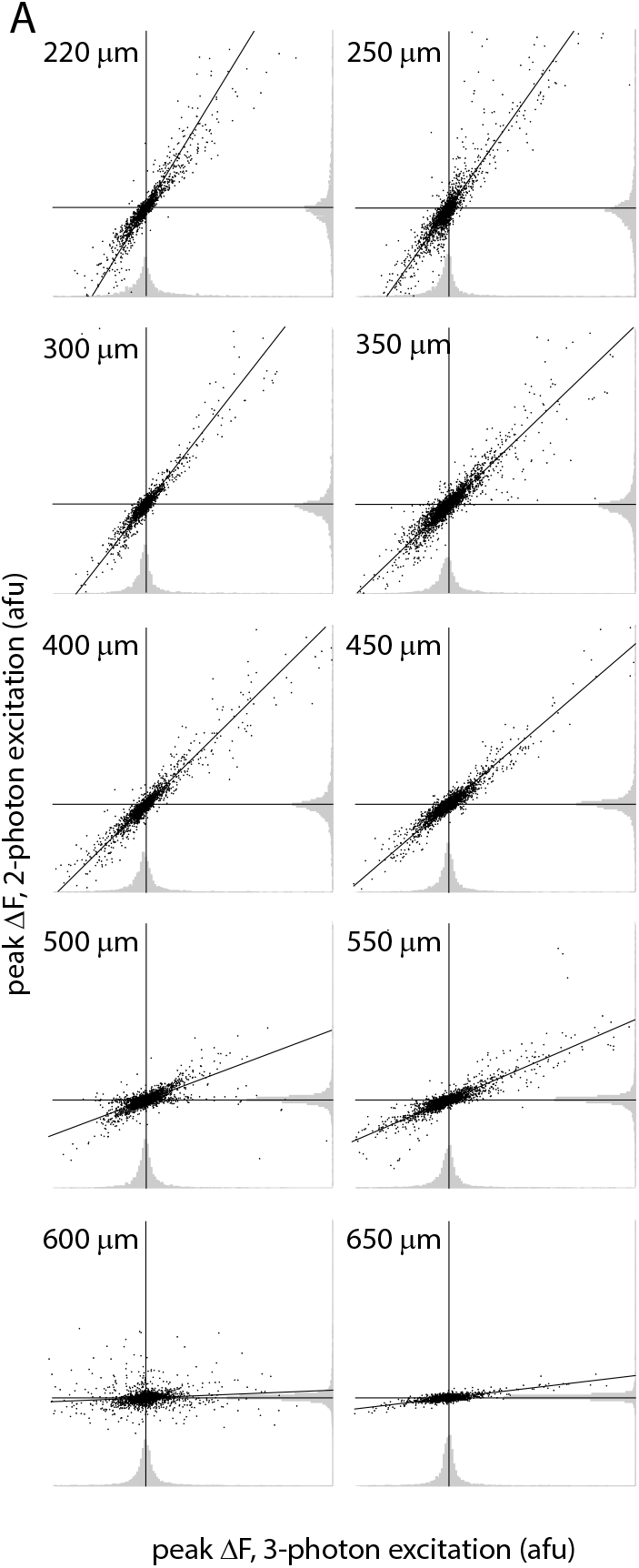
Fluorescence changes evoked by drifting gratings, measured with 2- and 3- photon excitation. Plots of 2- vs 3-photon changes in fluorescence evoked by drifting gratings. 10 plots illustrate results from 10 depths. x and y axes each display peak changes in fluorescence (ΔF) from −50 to +100 arbitrary fluorescence units. x axis: 3-photon ΔF. y axis: 2-photon ΔF. Each plot shows pooled results from many neurons, with each neuron contributing 24 data points (12 directions, 2 spatial frequencies). Each plot includes a line of best fit. Histograms display the distribution of data points on each plot; 2-photon distribution below each plot and 2-photon distribution to the right.

**Supplementary figure 4.**
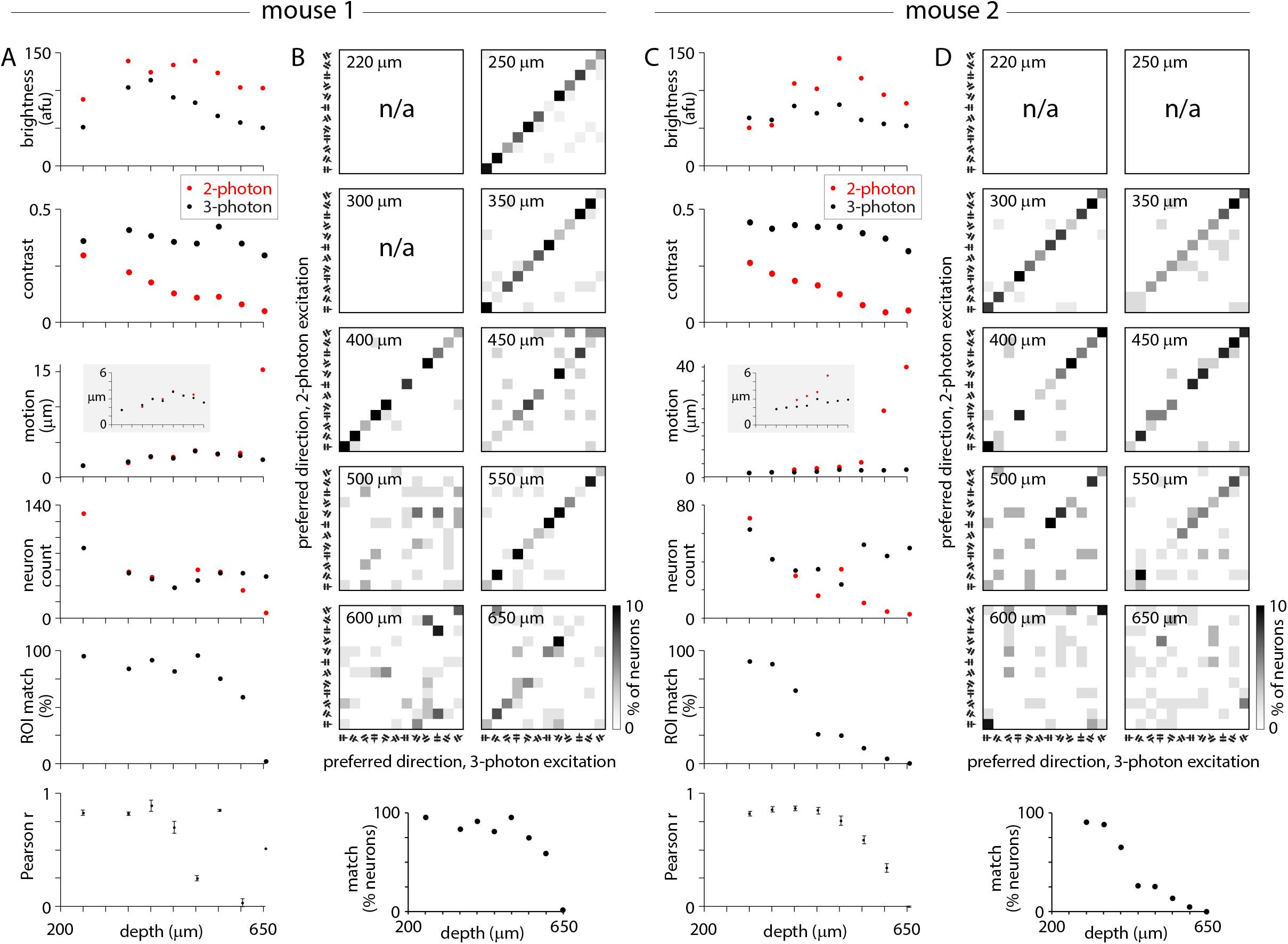
Depth-dependent changes in image quality and apparent direction preference differ between mice. (A) Image quality as a function of depth for a single mouse. Equivalent to plots in figure 3A-F. (B) Direction preference as a function of depth for the same mouse as panel A. Equivalent to plots in figure 3H & J. (D, E) Same plots for a different mouse. Insets in motion plots: same x axis, expanded y axis.

**Supplementary figure 5.**
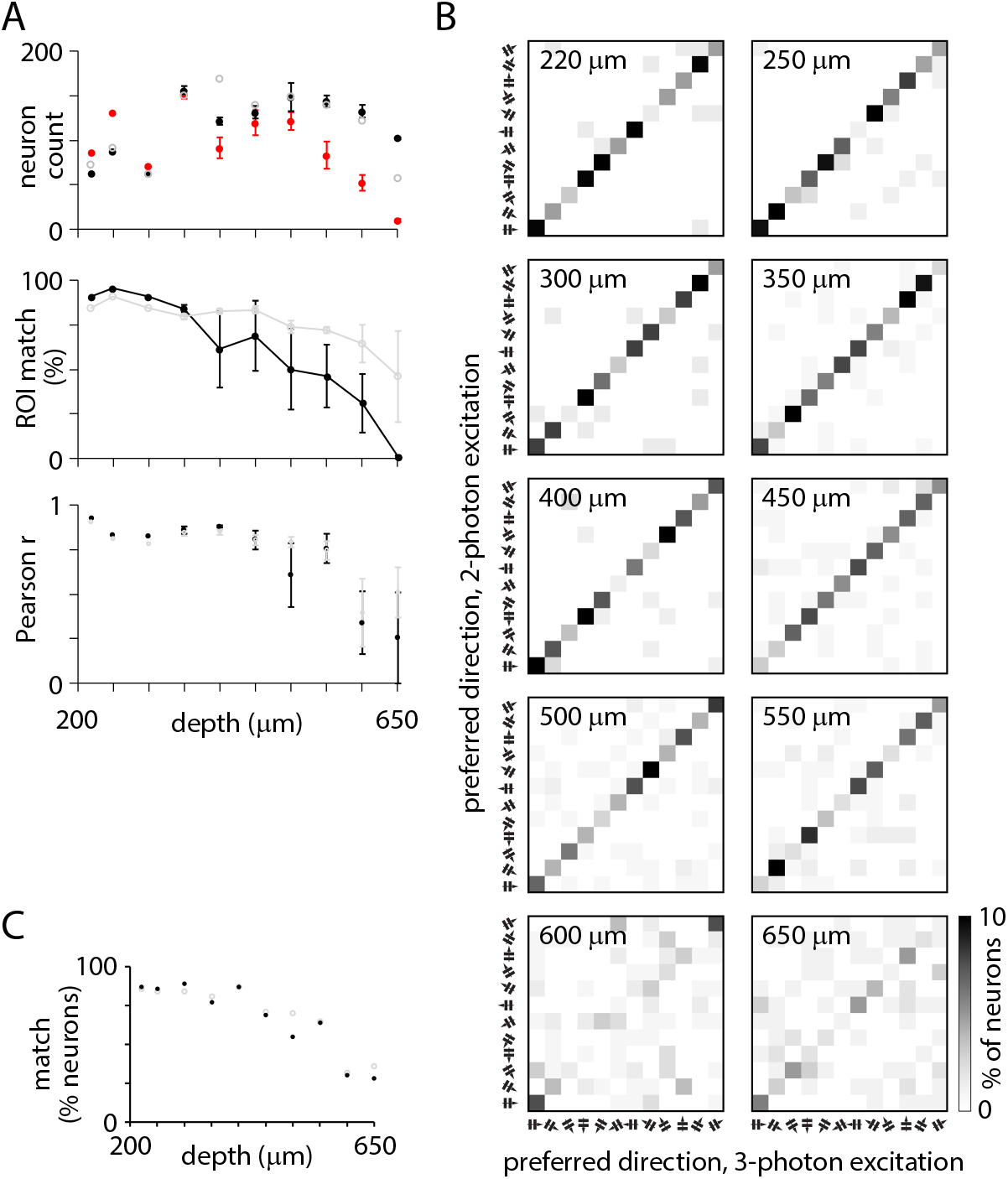
Comparison of 2- and 3-photon results using 3-photon motion correction. Correcting motion in our 2-photon images using estimates of motion from 3-photon images improved segmentation, but failed to recover accurate direction preference from 2-photon measurements. Image quality as function of depth (A) and preferred directions (B) after correction of 2-photon images with motion information from 3-photon images. Black and red results are duplicates of those in figure 4. Grey symbols indicate 2-photon results after motion correction with 3-photon motion estimate. Using motion estimated from the 3-photon images to motion-correct the corresponding deep-layer 2-photon images improved segmentation from 2-photon images, increasing cell count and % overlap, but there was little change in the Pearson correlation coefficient, the slope of the relationship between 2- and 3-photon fluorescence changes failed to recover, and the number of neurons with matched preferred direction remained low. Hence improved motion correction failed to enable extraction of accurate fluorescence results from 2-photon movies in deep locations. Presumably fluorescence emitted by GCaMP from deep-layer neurons after 2-photon excitation accurately reports direction preference. We expect the neuropil-subtraction routine to have subtracted the mean of the out-of-focus background, but the photon noise associated with this background was presumably sufficient to obscure the preferred direction of many neurons. The dominance of out-of-focus background may have been facilitated by the adjustment of laser illumination to maintain approximately the same mean fluorescence per image at each depth, resulting in weak excitation of GCaMP6s in deep-layer neurons.

**Supplementary figure 6.**
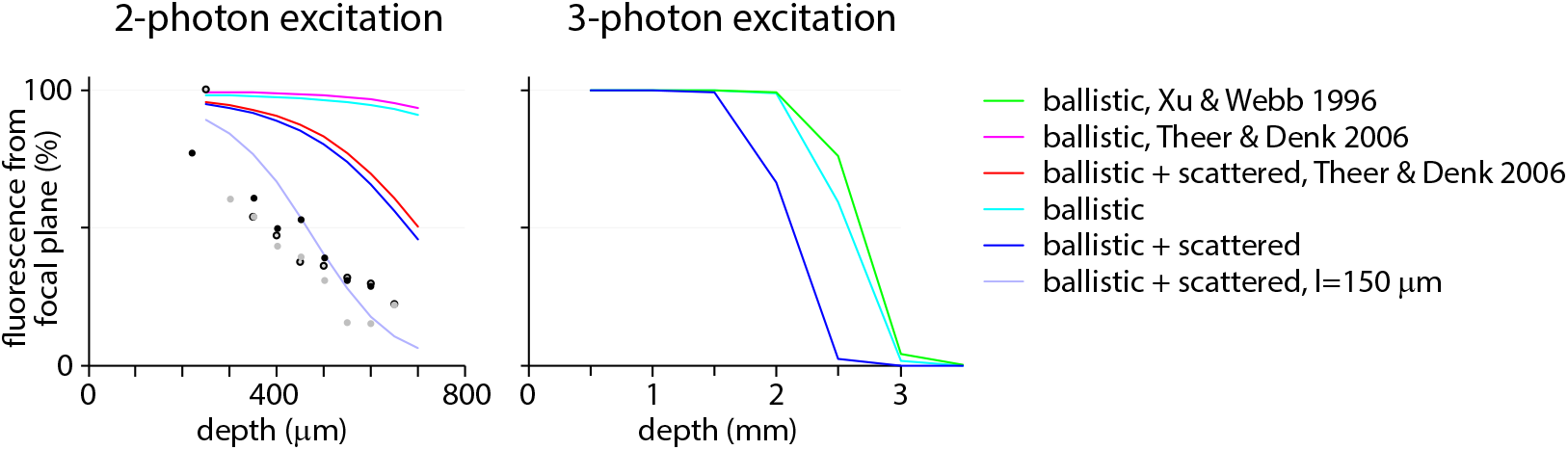
In-to out-of-focus fluorescence ratio. (A) Percentage of fluorescence originating from the focal plane, estimated using analytical expressions and plot as a function of the depth of the focal plane below the brain surface. Points, measurements from 3 experiments (black, grey, open symbols). Lines, calculated values using equation 7 of Xu & Webb, 1996 (green), equation 4 of Theer & Denk, 2006 (pink, red), and our equation 1 (blues). Lines were calculated with a scattering length constant of 200 μm except one line, calculated using our equation 1 and a scattering length constant of 150 μm.

**Supplementary figure 7.**
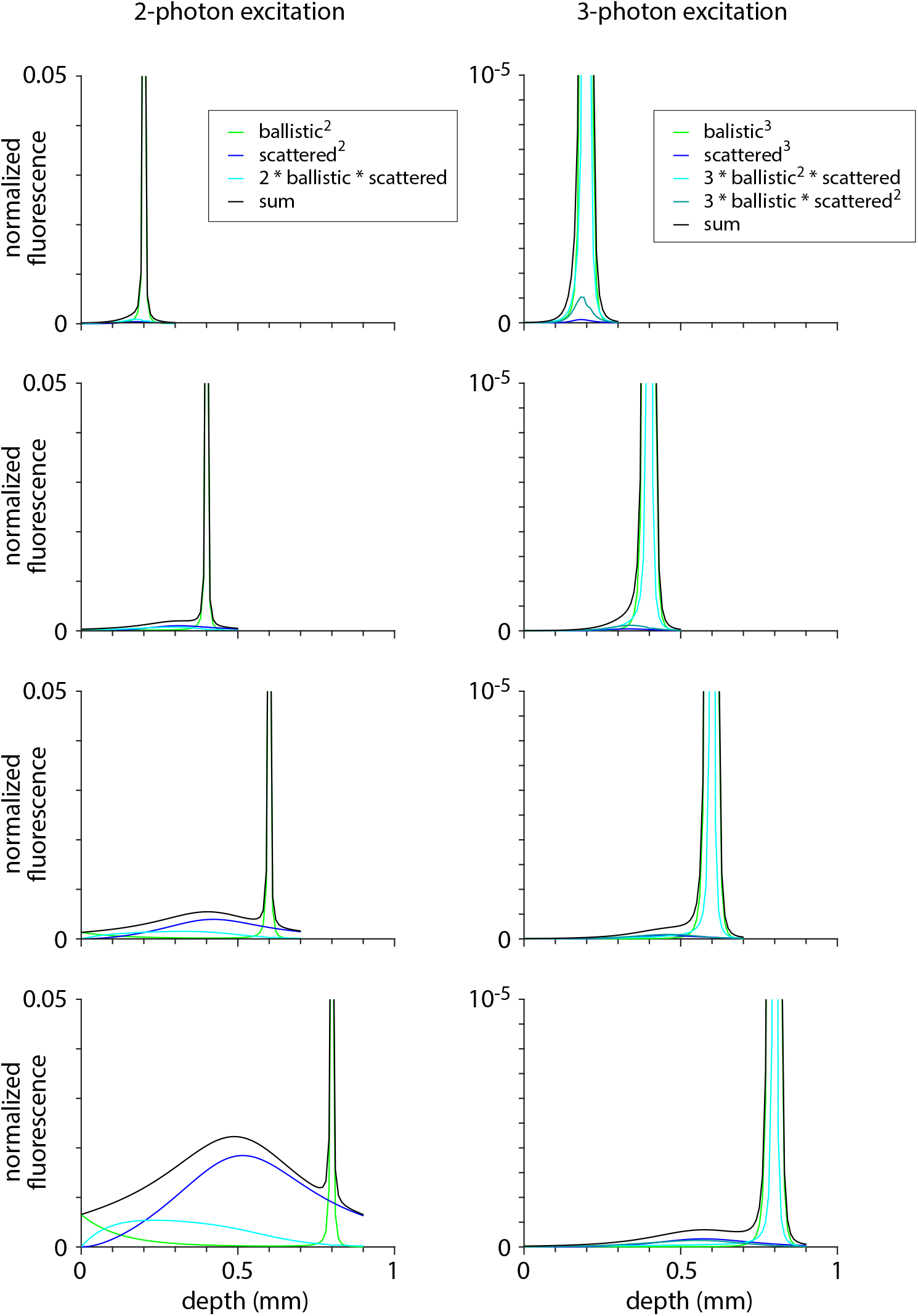
Estimated fluorescence from in-and out-of-focus planes. Plot illustrating the depth from which fluorescence originates (see Theer & Denk, 2006). Lines for 2-photon excitation were calculated for 900 nm illumination and scattering length 200 μm using equation 1. Lines for 3-photon excitation were calculated for 1300 nm illumination and scattering length 200 μm using an equivalent formulation. Colors indicate fluorescence from ballistic incident photons (light green), from scattered photons (dark blue) and from a mixture of ballistic and scattered photons (cyan and deep green). Black: the sum of all fluorescence sources (reproduced in figure 4B).

## REFERENCES

Akassoglou K, Merlini M, Rafalski VA, Real R, Liang L, Jin Y, Dougherty SE, De Paola V, Linden DJ, Misgeld T, Zheng B (2017) *In vivo* imaging of CNS injury and disease. J Neurosci 37(45), 10808–10816.

Bertie JE, Lan Z (1996) Infrared Intensities of Liquids XX: The Intensity of the OH Stretching Band of Liquid Water Revisited, and the Best Current Values of the Optical Constants of H_2_O(l) at 25°C between 15,000 and 1 cm^-1^. Applied Spectroscopy 50, 1047–1057.

Carrillo-Reid L, Yang W, Miller JK, Peterka DS, Yuste R (2017) Imaging and optically manipulating neuronal ensemble. Ann Rev Biophys 46, 271–293.

Curcio JA, Petty CC (1951) The near infrared absorption spectrum of liquid water. J Opt Soc Am 41(5), 302–304.

Daigle TL, Madisen L, Hage TA, Valley MT, Knoblich U, Larsen RS, Takeno MM, Huang L, Gu H, Larsen R, et al. (2018). A suite of transgenic driver and reporter mouse lines with enhanced brain-cell-type targeting and functionality. Cell 174, 465–480.

Dana H, Sun Y, Mohar B, Hulse B, Hasseman JP, Tsegaye G, Tsang A, Wong A, Patel R, Macklin JJ, Chen Y, Konnerth A, Jayaraman V, Looger LL, Schreiter ER, Svoboda K, Kim DS (2018) High-performance GFP-based calcium indicators for imaging activity in neuronal populations and microcompartments. bioRxiv 434589.

Denk W, Delaney KR, Gelperin A, Kleinfeld D, Strowbridge BW, Tank DW, Yuste R (1994) Anatomical and functional imaging of neurons using 2-photon laser scanning microscopy. J Neurosci Meth 54, 151–162.

Denk W, Svoboda K (1997) Photon upmanship: why multiphoton imaging is more than a gimmick. Neuron 18, 351–357.

de Vries SEJ, Lecoq J, Buice MA et al. (2018) A large-scale, standardized physiological survey reveals higher order coding throughout the mouse visual cortex. bioRxiv 359513.

Goldey GJ, Roumis DK, Glickfeld LL, Kerlin AM, Reid RC, Bonin V, Schafer DP, Andermann ML (2014) Removable cranial windows for long-term imaging in awake mice. Nat Protoc 9, 2515–2538.

Gorski JA, Talley T, Qiu M, Puelles L, Rubenstein JL, Jones KR (2002) Cortical excitatory neurons and glia, but not GABAergic neurons, are produced in the Emx1-expressing lineage. J Neurosci 22, 6309–6314.

Hale GM, Querry MR (1073) Optical constants of water in the 200-nm to 200μm wavelength region. Applied Optics 12(3), 555–563.

Harris JA, Hirokawa KE, Sorensen SA, Gu H, Mills M, Ng LL, Bohn P, Mortrud M, Ouellette B, Kidney J, Smith KA, Dang C, Sunkin S, Bernard A, Oh SW, Madisen L, Zeng H (2014) Anatomical characterization of Cre driver mice for neural circuit mapping and manipulation. Front Neural Circuits 8, 76.

Holtmaat A, Randall J, Cane M (2013) Optical imaging of structural and functional synaptic plasticity *in vivo*. Eur J Pharmacol 719, 128–136.

Horton NG, Wang K, Kobat D, Clark CG, Wise FW, Schaffer CB, Xu C (2013) *In vivo* three-photon microscopy of subcortical structures within an intact mouse brain. Nature Photonics 7, 205–209.

Ji N, Milkie DE, Betzig E (2010) Adaptive optics via pupil segmentation for high-resolution imaging in biological tissues. Nature Methods 7(2), 141–147.

Ji N, Sato TR, Betzig E (2012) Characterization and adaptive optical correction of aberrations during in vivo imaging in the mouse cortex. PNAS 190(1), 22–27.

Kobat D, Durst ME, Nishimura N, Wong AW, Schaffer CB, Xu C (2009) Deep tissue multiphoton microscopy using longer wavelength excitation. Optics Express 17(16), 13354–13364.

Kobat D, Horton NG, Xu C (2011) In vivo two-photon microscopy to 1.6-mm depth in mouse cortex. J Biomed Opt 16(10), 106014.

Leray A, Odin C, Huguet E, Amblard F, Le Grand Y (2007) Spatially distributed two-photon excitation fluorescence in scattering media: Experiments and time-resolved Monte Carlo simulations. Optics Communications 272, 269–278

McLean JW, Freeman JD, Walker RE (1998) Beam spread function with time dispersion. Applied Optics 37, 4701–4711.

Ouzounov DG, Wang T, Wang M, Feng DD, Horton NG, Cruz-Hernández JC, Cheng YT, Reimer J, Tolias AS, Nishimura N, Xu C (2017) In vivo three-photon imaging of activity of GCaMP6-labeled neurons deep in intact mouse brain. Nature Methods 14, 388–329.

Pachitariu M, Stringer C, Dipoppa M, Schröder M, Rossi LF, Dalgleish H, Carandini M, Harris KD (2017) Suite2p: beyond 10,000 neurons with standard two-photon microscopy. bioRxiv 061507.

Podgorski K, Ranganathan G (2016) Brain heating induced by near-infrared lasers during multiphoton microscopy. J Neurophysiol. 116, 1012–1023.

Sigler A, Murphy T (2010) In vivo 2-photon imaging of fine structure in the rodent brain before, during, and after stroke. Stroke 41, S117–S123.

Theer P, Denk W (2006) On the fundamental imaging-depth limit in two-photon microscopy. J Opt Soc Am A 23(12), 3139–3149.

Theer P, Hasan MT, Denk W (2003) Two-photon imaging to a depth of 1000 mm in living brains by use of a Ti:Al2O3 regenerative amplifier. Optics Letters 28(12), 1022–1024.

Wang T, Ouzounov DG, Wang M, Xu C (2017) Quantitative comparison of two-photon and three-photon activity imaging of GCaMP6s-labeled neurons *in vivo* in the mouse brain. Optics in the Life Sciences.

Xu C, Webb WW (1996) Measurement of two-photon excitation cross sections of molecular fluorophores with data from 690 to 1050 nm. J Opt Soc Am B 13, 481–491.

Yildirim M, Sugihara H, So PTC, Sur M (2019) Functional imaging of visual cortical layers and subplate in awake mice with optimized three photon microscopy. Nature Communications 10:177, 1–12.

Ying J, Liu F, Alfano RR (1999) Spatial distribution of two-photon-excited fluorescence in scattering media: erratum. Applied Optics 38(10), 2151.

